# Feed Composition and Antibiotic Supplements Modulate Digestive Enzyme Activities and Alter Gut Microbiome of Pacific White Shrimp

**DOI:** 10.64898/2026.01.07.698159

**Authors:** Alexandru S. Barcan, Eve Hughes, Trond M. Kortner, Brendan Robertson, Joseph L. Humble, Martin S. Llewellyn

**Affiliations:** SBOHVM, University of Glasgow, Graham Kerr Building, G12 8QQ; Department of Paraclinical Sciences, Faculty of Veterinary Medicine, Norwegian University of Life Sciences (NMBU), Ås, Norway; Institute for Ocean Engineering, Shenzhen International Graduate School, Tsinghua University, 518055 Shenzhen, China

**Keywords:** Aquafeed, Microbiota, Digestibility, Enzymes, Gentamicin, Guar, Shrimp, Soya, Feather meal

## Abstract

The transition from fishmeal to sustainable alternatives in aquaculture is essential, however the physiological and microbial impacts of alternative diets in shrimp remain poorly understood. Here, we examine how substituting fishmeal with plant-based proteins such as guar and soybean meals, the inclusion of feather meal, and the use of a commonly used antibiotic (gentamicin) influence digestive enzyme function, protein digestibility, and gut microbial assemblages in *Litopenaeus vannamei* (Pacific white shrimp). The guar-based diet notably altered gut microbiota composition and decreased leucine aminopeptidase activity while maintaining high protein digestibility (>90%). In contrast, the soya/feather diet caused greater disruption to enzyme activity and microbial communities, resulting in reduced digestibility (∼75%). The gent/guar diet showed comparable digestibility and microbial stability to the guar diet, with only minor shifts at the genus level. Although digestibility data for the acclimation diet were unavailable, these findings highlight diet-specific physiological and microbial responses to fishmeal substitutes. This emphasizes the need to consider dietary formulation, digestive function, and microbiome dynamics when developing sustainable aquafeeds for shrimp farming.

**Importance:** The Pacific white shrimp (*Litopenaeus vannamei*) is a cornerstone of global aquaculture, yet optimizing its diet remains challenging. Current shrimp farming heavily depends on fishmeal, an unsustainable protein source, and antibiotic use to maintain shrimp health raises concerns about antimicrobial resistance and environmental impacts. This study highlights how alternative dietary formulations, including plant-based proteins and antibiotic supplements, influence shrimp digestive physiology and reshape gut microbiomes. Understanding these interactions is crucial to developing feed formulations that support robust shrimp growth and health without excessive reliance on antibiotics or fishmeal. By demonstrating how specific dietary ingredients affect both shrimp digestion and beneficial gut bacteria, this research provides valuable insights that can inform sustainable and responsible shrimp aquaculture practices globally.

## 1. Introduction

Aquaculture has become a cornerstone of global food security, with the production of species such as *Litopenaeus vannamei* playing a pivotal role in meeting the increasing demand for sustainable seafood (1). Nonetheless, optimizing the health and performance of farmed shrimp across diverse dietary and environmental conditions remains a significant challenge (2, 3). Digestive efficiency and gut microbiome composition are integral to shrimp growth (4), nutrient utilization (5, 6) and resilience against disease (4, 7), however our understanding of how dietary interventions modulate these factors is still limited. The shrimp aquaculture industry, valued at $40 billion in 2021, depends heavily on species like *L. vannamei* and *Penaeus monodon* (8). Among these, *L. vannamei* has gained global prominence due to its comparative resilience to diseases such as white spot syndrome (9), rapid growth rate (10), ability to tolerate lower water quality than other species (11), and its high culinary appeal. In their natural habitat, *L. vannamei* are opportunistic omnivores, feeding on fish, crustaceans, algae, and insects (12, 13). In contrast, aquaculture systems, particularly in semi-intensive pond systems, rely heavily on formulated feeds to meet their nutritional needs (1, 5, 14). These feeds often contain 20-50% fishmeal, a protein-rich ingredient derived from wild-caught fish (14, 15). Given the declining availability of fishmeal due to overfishing and climate change, there is an urgent need to develop alternative protein sources to meet the growing demand for aquafeeds (16, 17). Recent advancements in shrimp aquaculture have introduced innovative dietary and management solutions to address these challenges (15, 18, 19). Functional feeds incorporating probiotics (20), prebiotics (21), and postbiotics (22, 23) have demonstrated potential in improving shrimp health and growth. Additionally, plant-based protein sources such as guar meal and soybean meal are being explored as sustainable alternatives to fishmeal (18). Guar meal, derived from *Cyamopsis tetragonoloba*, is a cost-effective protein source with the potential to partially replace fishmeal when appropriately processed to reduce its anti-nutritional factors, such as guar gum (24). Its high fiber content, however, requires precise formulation to ensure nutrient digestibility (25). Research on rainbow trout diets demonstrated that a proprietary guar protein concentrate could successfully replace traditional protein sources without negatively affecting growth performance or gut health, underscoring its viability in aquaculture (25). Likewise, yeast-fermented guar meal combined with copra meal was found to substitute up to 75% of fishmeal in Nile tilapia diets, with no adverse impacts on growth, suggesting that fermentation is an effective method for enhancing the nutritional quality of guar (26). Further highlighting its potential, guar meal, when properly processed, has been shown to reduce costs and enhance sustainability in aquafeed production, positioning it as an appealing alternative to reduce dependence on traditional fishmeal (27). Feather meal, a byproduct of the poultry industry (28), is a protein-rich alternative increasingly used in aquaculture diets. Despite its high crude protein content (up to 85%), its use in shrimp diets is limited by the low digestibility of keratin, the primary protein in feathers (29). Hydrolyzed feather meal, processed to improve digestibility, has been successfully incorporated into diets for species like tilapia (30) and catfish (31), demonstrating its feasibility as a protein substitute. In *Litopenaeus vannamei*, low to moderate levels of hydrolyzed feather meal effectively replace fishmeal, though higher inclusion levels may reduce protein efficiency due to indigestible keratin (15, 32).

Similarly, soybean meal (SBM), valued for its high protein content and availability, has been widely utilized in shrimp diets (18). Contrary to its benefits, the inclusion of SBM must be carefully managed to avoid negative impacts from anti-nutritional factors, which can be mitigated through fermentation or enzymatic treatments (33). Recent research highlights the potential of SBM as a sustainable alternative to fishmeal in shrimp diets (34–36). Fermented SBM (FSBM) has been shown to enhance growth performance, feed utilization, and immune responses when included at moderate levels, with one study reporting optimal outcomes at 20% replacement of fishmeal in juvenile *Litopenaeus vannamei* diets, while higher levels led to reduced performance (35). Additionally, replacing up to 40% of fishmeal with SBM or FSBM maintained growth and immune parameters, demonstrating its viability as a plant-based protein source (34). Understanding the digestive physiology of shrimp is crucial for optimizing the incorporation of alternative protein sources into aquafeeds. The shrimp digestive system has specialized compartments, each with specific functions that are influenced by pH and environmental temperature (37). Foregut (stomach) activity involves mechanical and chemical breakdown of food, typically under acidic conditions (pH ∼5) (37, 38). Midgut (hepatopancreas) function is central to enzyme production, such as amylase, leucine aminopeptidase, trypsin, and nutrient absorption, operating at a neutral to slightly alkaline pH of 7-8 (39, 40). Finally, the hindgut (intestine) helps absorb remaining nutrients and water while excreting waste, usually maintaining an alkaline pH of 8-9 (41). The activity within these compartments is closely linked to the shrimp’s environment, with digestive processes optimized for water temperatures between 28-32°C (27, 42). Antibiotics are sometimes included in shrimp diets to manage bacterial infections and improve survival rates (43–45). However, their overuse has led to the emergence of antibiotic-resistant pathogens, significantly reducing their effectiveness and raising serious concerns about antimicrobial resistance and environmental impacts (46). Consequently, combining functional feed ingredients and optimized formulations can enhance shrimp health and reduce dependency on antibiotics, addressing the growing concern of antimicrobial resistance and environmental impacts (47, 48). In response to these issues, our study focused on evaluating three dietary strategies: a guar meal-based diet, a soybean/feather meal mix (soya/feather), and a gentamicin-treated guar diet (gent/guar). We investigated their effects on digestive enzyme activity, nutrient digestibility, and gut microbiome composition across different gut compartments of *L. vannamei.* Through enzymatic assays and microbial profiling, our findings provide novel insights into how distinct dietary inputs influence shrimp physiology and gut microbial communities, offering actionable strategies to reduce fishmeal dependence and promote resilient, eco-friendly shrimp aquaculture.

## 2. Materials and methods

### 2.1. Shrimp acclimation

Forty-four captive-bred *Litopenaeus vannamei* shrimp were purchased from RAS Scotland and housed in five 40 L tanks. Water parameters, including pH, temperature, ammonia, and nitrate concentrations, were monitored daily and adjusted as needed to maintain optimal conditions. A 50% water change was performed each day to ensure water quality. Shrimp were acclimated over one week and fed the acclimation diet ad libitum. Following the acclimation period, four shrimp were randomly selected and euthanized by chilling to 2°C in an ice slurry. The hepatopancreas, stomach, and midgut were dissected, with tissues stored at -80°C for subsequent microbiome analysis. A portion of the hepatopancreas was reserved for enzyme extraction.

### 2.2. Diet formulation for feeding trials

Two experimental rainbow trout diets were formulated and produced by extrusion at Skretting Aquaculture Innovation (Stavanger, Norway). Although standard shrimp feeds generally have lower lipid content, we opted to use a trout-based pellet matrix because it offers a well-characterized, consistent nutrient profile. This uniformity allowed us to compare the effects of different diet formulations, each incorporating distinct protein sources such as guar meal or soya/feather meal, without introducing confounding batch-to-batch differences in feed composition. We recognize that trout pellets differ nutritionally from commercial shrimp rations, however, this controlled approach enabled us to generate mechanistic insights that can inform future shrimp-specific feed formulations.

The guar-based diet included 20% guar meal, 18% wheat gluten, and 13.53% wheat, while the soya/feather diet was formulated with 21.53% soybean meal, feather meal 20%, 10% wheat gluten, and 11.08% wheat. The formulation parameters allowed variation also in some other protein rich ingredients than those of most interest in the present studies, i.e. ingredients with high digestibility such as wheat gluten and soy protein concentrate. Wheat meal was used to balance the carbohydrate level of the diets. Both diets incorporated yttrium as an inert marker to facilitate digestibility analysis. Detailed compositions of these diets are provided in table 1. A third diet, referred to as the antibiotic supplement diet, was prepared by dissolving 12 mL of gentamicin in distilled water and thoroughly mixing it into the guar-based formulation. The final gentamicin concentration in the diet was 25 mg/kg. As controls, 12 mL of distilled water was added to both the guar-based and soya/feather-based feeds. All feed formulations were coated with cod liver oil and dried in a dehydrator at 60°C for 24 hours until solid. The cod liver oil was applied as a top coat to all diets. In the gentamicin-supplemented formulation, this step was essential to limit leaching of gentamicin into the water, as the antibiotic is water-soluble. During the one-week pretrial period, shrimp were fed a commercially available maintenance diet for *L. vannamei* supplied by RAS Scotland.

**Table 1.**
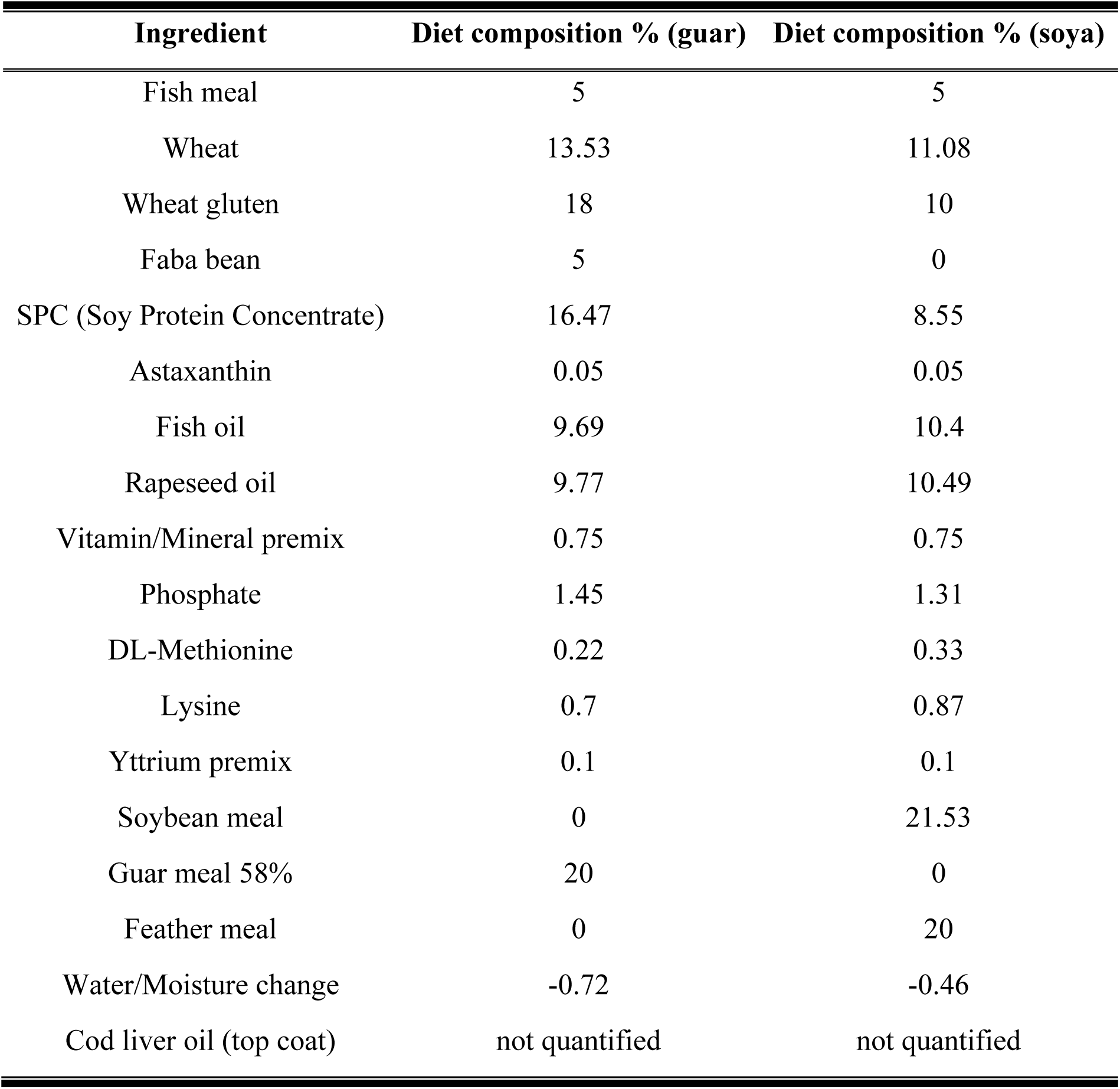
Ingredient composition of trout pellet diets with guar, soya and feather meal.

| Ingredient | Diet composition % (guar) | Diet composition % (soya) |
| --- | --- | --- |
| Fish meal | 5 | 5 |
| Wheat | 13.53 | 11.08 |
| Wheat gluten | 18 | 10 |
| Faba bean | 5 | 0 |
| SPC (Soy Protein Concentrate) | 16.47 | 8.55 |
| Astaxanthin | 0.05 | 0.05 |
| Fish oil | 9.69 | 10.4 |
| Rapeseed oil | 9.77 | 10.49 |
| Vitamin/Mineral premix | 0.75 | 0.75 |
| Phosphate | 1.45 | 1.31 |
| DL-Methionine | 0.22 | 0.33 |
| Lysine | 0.7 | 0.87 |
| Yttrium premix | 0.1 | 0.1 |
| Soybean meal | 0 | 21.53 |
| Guar meal 58% | 20 | 0 |
| Feather meal | 0 | 20 |
| Water/Moisture change | -0.72 | -0.46 |
| Cod liver oil (top coat) | not quantified | not quantified |

### 2.3. Feeding trials

Four shrimp were randomly assigned to each of the three experimental diets, while the remaining shrimp continued the acclimation diet. Treatment shrimps were weighed prior to the trial. On the first day, the treatment groups were fed a diet comprising 50% acclimation feed and 50% treatment diet. From the second day onward, the shrimp were fed the treatment diet exclusively, three times daily at 10:00 am, 12:00 pm, and 15:00 pm. Feeding times for the three groups were staggered by five minutes. Uneaten feed was removed after 30 minutes using a Pasteur pipette. Faeces were collected from treatment tanks at 15-minute intervals for one hour after feeding, rinsed with distilled water, and pooled. Samples were stored at −4°C for later analysis. Faeces were not collected after the 10:00am feeding if shrimp in the tank had moulted overnight. Faeces collection was carried out over six days. At the end of the feeding trial, the shrimp were euthanized by chilling in an ice slurry. The hepatopancreas, stomach, and midgut were dissected, with tissues stored at −80°C for microbiome analysis. A portion of the hepatopancreas was used for enzyme extraction. The feeding trial spanned three weeks, with four shrimp assigned to each experimental diet (n = 4 × 3 = 12).

### 2.4. Enzyme activity assays

The activities of total protease, leucine aminopeptidase, amylase, and trypsin were measured using hepatopancreas enzyme extracts. To analyze the effects of different diets, enzyme activities from shrimp fed the acclimation diet were compared with those from shrimp on the three treatment diets. Data normality was assessed using the Shapiro-Wilk test. Differences in enzyme activity across diets were evaluated using a one-way ANOVA, followed by Tukey’s post-hoc test to identify significant pairwise differences.

### 2.5. Enzyme extraction

Tissue samples were combined with chilled distilled water at a ratio of 6 mL water per gram of tissue. The hepatopancreas homogenates were processed using a stick homogenizer for 2 minutes. The homogenate was centrifuged at 12,000g for 20 minutes at 4°C, and the supernatant was aliquoted. The crude enzyme extract was stored at -80°C for later analysis.

### 2.6. Enzyme activity analysis

Enzyme activity from hepatopancreas extracts was analyzed to assess the functional capacity of digestive enzymes under different dietary treatments. Enzyme activity was defined as the amount of enzyme required to catalyze the conversion of one mole of substrate per minute under specified conditions (49). All assays were performed in triplicate under standardized conditions of 40°C and pH 8.

### 2.7. Total protease activity

Total protease activity was assessed using an adapted method from Cupp-Enyard (2008) (50). The substrate consisted of 0.65% (w/v) casein dissolved in 50 mM potassium phosphate buffer. The solution was heated to 80°C until fully dissolved, cooled to room temperature, and pH-adjusted using HCl or NaOH. For each reaction, 0.2 mL of enzyme extract was mixed with 5 mL of substrate solution and incubated for 10 minutes. Blank reactions were prepared by replacing the enzyme with 0.2 mL of distilled water.

Reactions were terminated by the addition of 5 mL 110 mM trichloroacetic acid (TCA), filtered through a 0.45 µm polyethersulfone syringe, and 2ml of filtrate was combined with 5 mL 500 mM sodium carbonate, and 1 mL 0.5 mM Folin’s phenol reagent, the samples were incubated at 37C for 30 mins, and analyzed spectrophotometrically for absorbance was recorded at 660 nm. A standard curve was prepared using L-tyrosine to quantify tyrosine release. Enzyme activity (U/mL) was calculated using the formula:

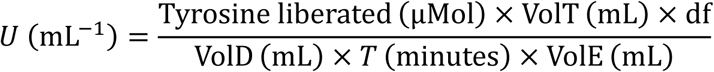

where VolT represents the total reaction volume, VolE the enzyme volume, VolD the detection volume, T the reaction time, and df the dilution factor.

### 2.8. Trypsin activity

The method for measuring trypsin activity was adapted from Erlanger et al. (1961) (51). The substrate, 11.5 mM BAPNA, was prepared in dimethyl sulfoxide (DMSO), protected from light, and stored at -4°C. A buffer solution containing 50 mM calcium chloride and 500 mM TRIS was prepared in distilled water, and 20 mL of this buffer was combined with 8.7 mL of the BAPNA solution. The substrate solution was adjusted to the desired pH using HCl or NaOH and diluted to 100 mL with distilled water. Reactions were initiated by adding enzyme extract to the substrate solution, with blanks prepared using distilled water in place of the enzyme. After 30 minutes of incubation at the specified temperature, reactions were terminated with 0.5 M HCl. The assay mixture was filtered through a 0.45 µm polyethersulfone syringe filter, and the absorbance of the filtrate was measured at 410 nm using a spectrophotometer. Enzyme activity was calculated using the following equation (49):

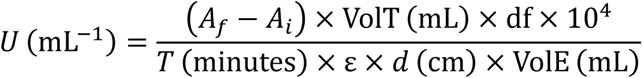

Where A_f_ represents the final absorbance value, A_I_ represents the initial absorbance value, VolT represents the total reaction volume, df represents the dilution factor, T represents the reaction time, ɛ represents the extinction coefficient of the substrate, d represents the optical path, and VolE represents the volume of enzyme used.

### 2.9. Leucine aminopeptidase activity

Leucine aminopeptidase activity was measured using a method adapted from the trypsin assay. The substrate, 11.5 mM LPNA, replaced BAPNA in the reaction mixture. The substrate solution was prepared in a similar manner, with the pH adjusted using HCl or NaOH and stored appropriately. Reactions were conducted in triplicate. The same calculation method used for trypsin activity was applied to determine leucine aminopeptidase activity, allowing for consistent comparison of enzyme kinetics across conditions.

### 2.10. Amylase activity

Amylase activity was measured using a method adapted from Bernfeld (1955) (52). The color reagent was prepared by dissolving 1496 mg/mL potassium sodium tartrate tetrahydrate in 2 M NaOH and combining this with a 21.9 mg/mL solution of 3,5-dinitrosalicylic acid dissolved in distilled water. The final color reagent consisted of 12 mL distilled water, 8 mL of the potassium sodium tartrate solution, and 20 mL of the dinitrosalicylic acid solution, mixed and heated to 80°C. The substrate solution was prepared by dissolving 2.4 mg/mL sodium phosphate and 0.39 mg/mL sodium chloride in distilled water. Potato starch (10 mg/mL) was added, and the mixture was boiled to ensure complete dissolution. The pH was adjusted to the desired value with HCl or NaOH. A standard curve was generated using a 0.2% (w/v) maltose solution. To initiate the assay, 0.2 mL of enzyme extract was added to 1 mL of substrate solution, followed by 1 mL of color reagent. Reactions were incubated in a boiling water bath for 15 minutes and subsequently cooled on ice. Blank reactions were prepared by adding 0.2 mL of enzyme extract after incubation. After the addition of 9 mL of distilled water, absorbance was measured at 540 nm using a spectrophotometer. Maltose released during the reaction was quantified using the standard curve, and amylase activity was calculated as:

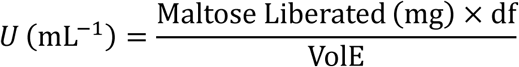

Where df represents dilution factor and VolE represents the volume of enzyme. The mean of enzyme activity for each temperature and pH was calculated.

### 2.11. Apparent digestibility

Faecal samples were homogenized for 2 minutes using ceramic sphere beads in a bead basher and subsequently freeze-dried. Yttrium concentration, used as an inert marker, was measured in both feed and faecal samples to assess digestibility. Analyses were conducted by the Molema lab (University of Glasgow). Protein concentrations in feed and faecal samples were determined using the Kjeldahl method, performed by Campden BRI. Apparent digestibility (AD) was calculated for each faecal sample using the following equation:

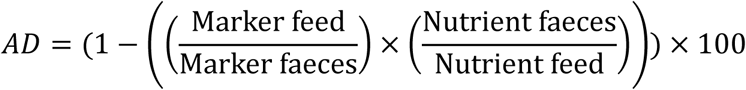

where “marker” refers to the yttrium concentration (mg/kg) and “nutrient” refers to the protein concentration.

Differences in apparent digestibility among the shrimp feeds were analyzed statistically. Data normality was tested using the Shapiro-Wilk test. Group differences were evaluated using ANOVA, followed by Tukey’s post-hoc test for pairwise comparisons.

### 2.12. Microbiome analysis

DNA extraction and library preparation for 16S rRNA sequencing were carried out using protocols based on established protocols (53, 54). Samples from the stomach, hepatopancreas, and midgut were processed using the QIAmp PowerFecal Pro DNA Kit according to the manufacturer’s instructions. DNA concentration was measured with a Qubit High Sensitivity Kit. Amplification of the V1 region of the 16S rRNA gene was performed using primers 27F (5’-AGAGTTTGATCMTGGCTCAG-3’) and 338R (5’-TGCTGCCTCCCGTAGGAGT-3’) under the following conditions: 95°C for 10 minutes (initial denaturation), 25 cycles of 95°C for 30 seconds, 55°C for 30 seconds, and 72°C for 30 seconds, and a final elongation at 72°C for 10 minutes. Selection of the V1 region minimized salmon DNA contamination, enhancing the accuracy of microbial community profiling (55, 56).

A two-step PCR approach was used to prepare samples for sequencing. The first PCR added internal barcodes, and products were verified on a 1% agarose gel. The second PCR introduced external barcodes, and the resulting products were again checked by agarose electrophoresis. Cleanup of PCR products was performed using the Biolabs Monarch Kit with a bead-to-sample ratio of 0.6. DNA concentration was confirmed using a Nanodrop spectrophotometer. Low initial DNA yields necessitated a second bead cleanup to further concentrate the DNA. The purified products were pooled at a final concentration of 10 nM and sequenced on a NovaSeq 6000 platform.

### 2.13. 16S rRNA sequencing data processing and visualization

Genus-level bacterial community data were analyzed based on gut segment location (stomach, midgut, hindgut) and dietary treatments (acclimation, antibiotic supplement, guar-based, soya/feather diets). Abundance values were aggregated across samples, and the top 25 genera were selected for detailed analysis. Sample names were parsed to associate data with gut compartments and dietary treatments, enabling compartment-based comparisons. Shannon diversity indices were calculated for each sample based on genus-level relative abundances to assess alpha diversity. Statistical analyses, including ANOVA and Tukey’s HSD tests, were performed to evaluate differences in Shannon diversity across gut segments and dietary treatments. Heatmaps and alpha diversity visualizations were created using Python’s Matplotlib (57) and Seaborn libraries (58). Non-metric multidimensional scaling (NMDS) plots were generated in R using the ggplot2 package (59). Differential abundance analysis was conducted using the DESeq2 package in R (60). Count data from 16S rRNA sequencing were grouped by gut segment and dietary treatment. Genera with low counts (≤10) were excluded from the analysis. Pairwise comparisons of genus abundances within each gut compartment were performed using DESeq2’s Wald test, with significant differences identified at an adjusted p-value (padj) < 0.05. Visualizations of differential abundance included a heatmap and bar plots, both generated in R using ggplot2 (59).

## 3. Results

### 3.1. Effects of dietary treatments on digestive enzyme activity

The enzyme activity of leucine aminopeptidase varied significantly among the different dietary treatments. The acclimation diet yielded the highest enzyme activity. Amylase activity was highest in the acclimation group, decreasing significantly in shrimp fed the guar-based and soya/feather-based diets. Leucine aminopeptidase activity remained consistently low across all diets. Protease activity showed substantial variation, peaking in the acclimation group, while trypsin activity was negligible across all treatments.

Statistical analysis ANOVA confirmed significant differences in leucine aminopeptidase activity between diets (p = 1.8e-05). Post-hoc Tukey tests further highlighted specific pairwise differences. The acclimation diet exhibited significantly higher activity compared to the gentamicin (p = 0.0018), guar (p = 0.0000116), and soya/feather (p = 0.0001354) diets. Among the experimental diets, the gentamicin and guar diets also differed significantly (p = 0.0004227), as did the guar and soya/feather diets (p = 0.0030950). However, no significant difference was detected between the gentamicin and soya/feather diets. For other enzymes, ANOVA revealed no significant differences among dietary treatments for trypsin (p = 0.173), protease (p = 0.399), and amylase (p = 0.167).

**Fig. 1.**
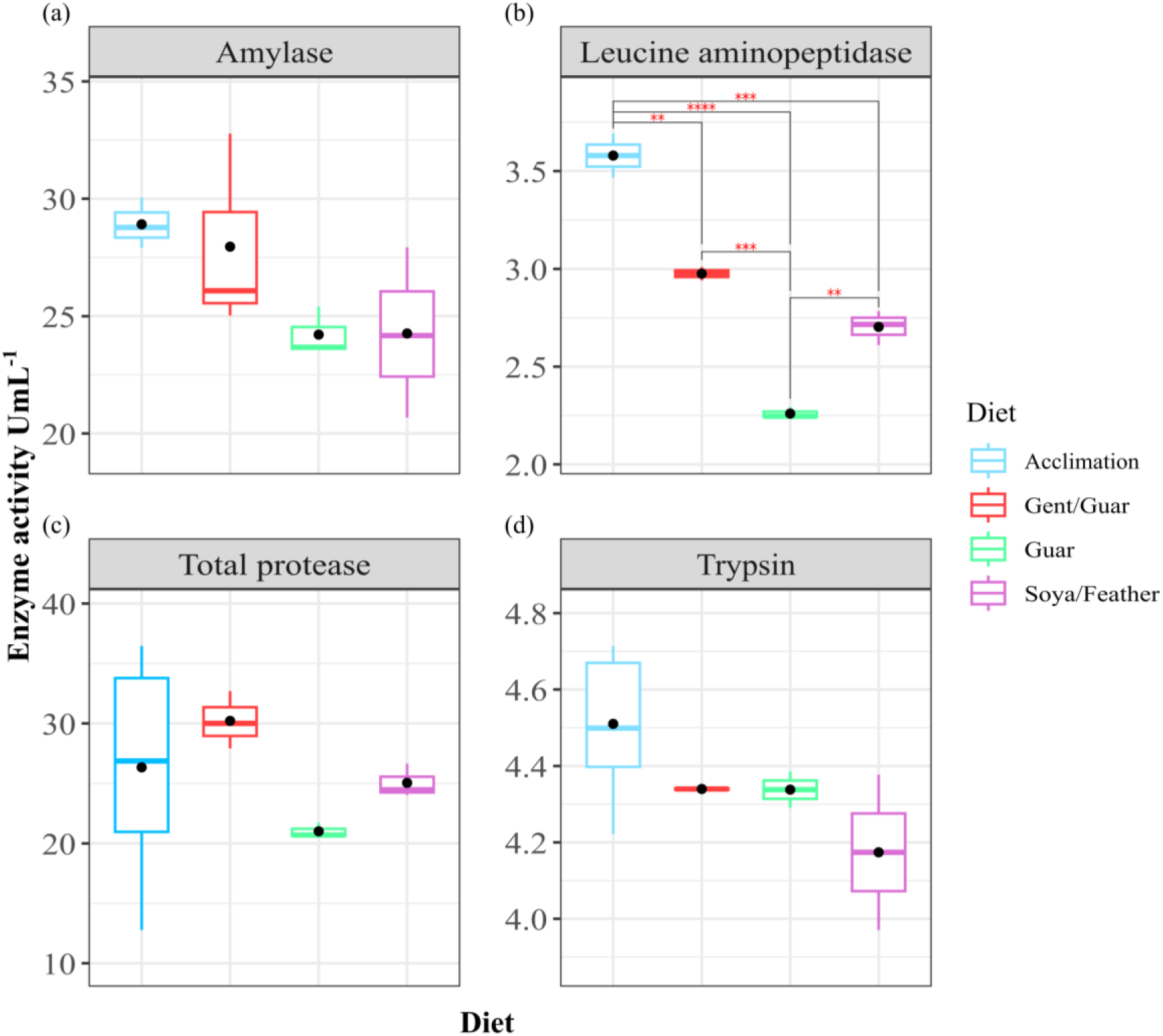
Comparative digestive enzyme activity across different dietary treatments. Boxplots depict the activity (U/mL) of amylase, leucine aminopeptidase, total protease, and trypsin in shrimp fed acclimation, gentamicin-supplemented, guar-based, and soya/feather-based diets.

### 3.2. Apparent digestibility of diets

The Apparent digestibility coefficients (ADCs) for dietary protein were measured across the experimental diets. As result, the ADCs for dietary protein differed significantly among the experimental diets (ANOVA, p = 8.66 × 10⁻⁶). Shrimp fed the gentamicin-supplemented diet exhibited the highest digestibility with a mean ADC of 92.69%, followed by the guar-based diet with a mean ADC of 90.88%. In contrast, the soya/feather-based diet had the lowest digestibility with a mean ADC of 72.68%. Post hoc Tukey tests confirmed that the gent/guar diet had higher digestibility than the soya/feather diet (p = 2.1 × 10⁻⁵), and the guar diet also showed greater digestibility than the soya/feather diet (p = 1.18 × 10⁻⁵). No significant difference in digestibility was observed between the gent/guar and guar diets.

**Fig. 2.**
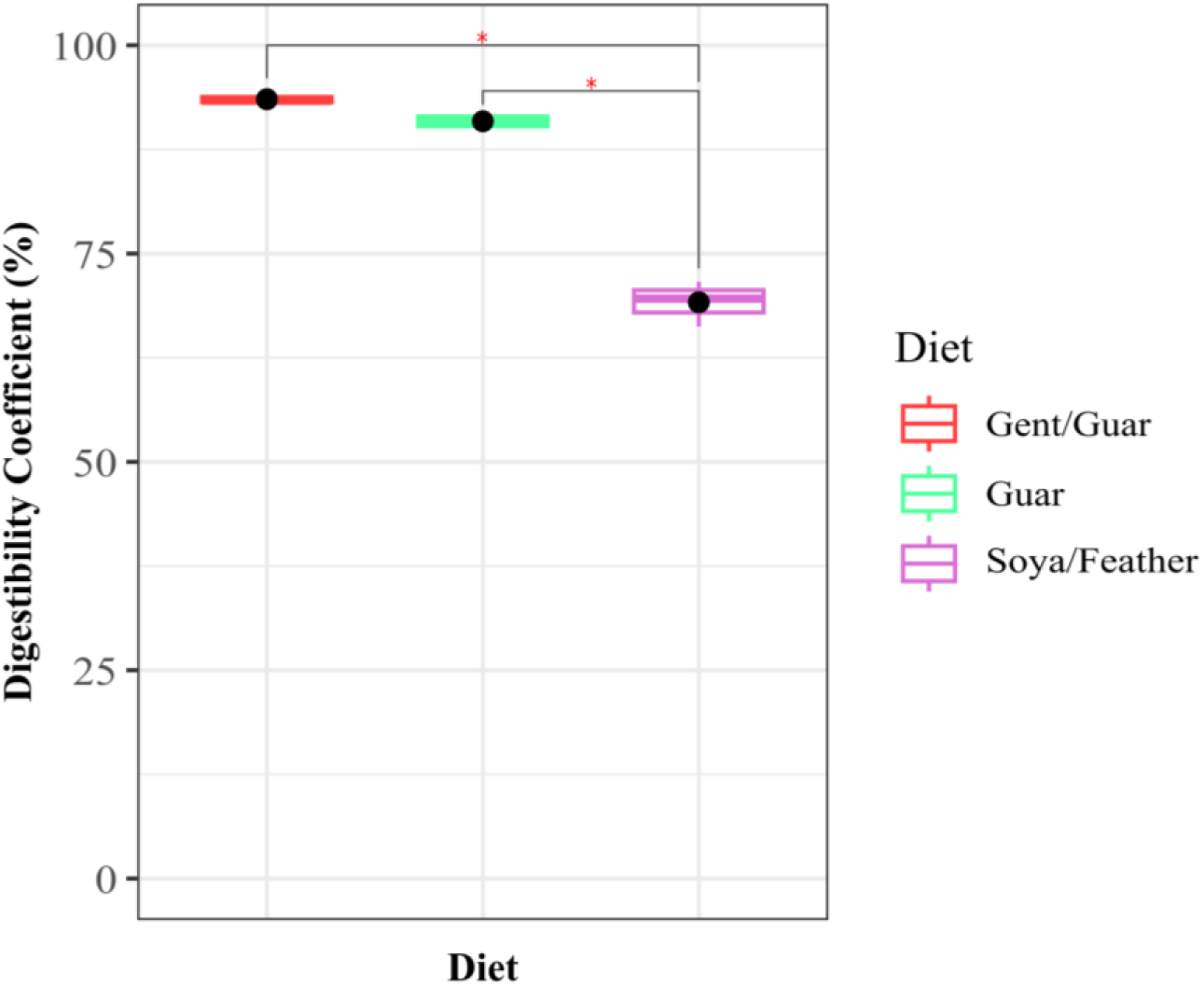
Apparent digestibility coefficients (ADCs) of protein in Litopenaeus vannamei fed different diets. Boxplots represent ADCs for shrimp fed gentamicin-supplemented, guar-based, and soya/feather-based diets. Shrimp fed the gentamicin and guar diets exhibited similarly high ADCs (above 90%), denoted by the letter “a.” In contrast, shrimp fed the soy-based diet had significantly lower ADCs (around 75%), denoted by the letter “b.” Statistical differences between diets were determined using ANOVA and Tukey’s post-hoc test (p < 0.05).

### 3.3. Microbial composition and diversity across gut compartments in *Litopenaeus vannamei* under different dietary treatments

We calculated the percentages of microbial reads relative to the total reads within each compartment to better understand the distribution of abundant genera across diets. In Figure 3a, we note a substantial presence of *Vibrio* across nearly all samples, indicating its wide distribution throughout the gut segments and dietary treatments.

The percentages of *Vibrio* populations in the guar diet were 12.35% in the stomach, 17.32% in the midgut, and 19.63% in the hindgut. In the gent/guar diet, these percentages increased to 18.94% in the stomach, 22.08% in the midgut, and 37.01% in the hindgut, suggesting that the addition of gentamicin amplifies *Vibrio* populations, particularly in the midgut and hindgut. For the soya/feather diet, the percentages were 14.34% in the stomach, 16.95% in the midgut, and 50.11% in the hindgut. Additionally, other genera showed notable trends. For example, *Demequina* accounted for 16.70% of the reads in the stomach under the soya/feather diet, which is higher compared to other diets such as guar (14.92%) and gent/guar (15.92%). Similarly, in the midgut, its abundance was 13.70%, slightly higher than guar (12.62%) and gent/guar (11.48%). However, in the hindgut, its abundance (7.31%) was lower than that observed in the guar diet (10.63%).

*Ruegeria* was most abundant in the stomach under the soya/feather diet (18.60%) but had lower percentages in the midgut (8.09%) and hindgut (10.22%). *Photobacterium* showed a more even distribution across diets and compartments, with the highest percentage in the hindgut under the acclimation diet (27.45%).

These percentages reveal distinct compartment-specific and diet-associated dynamics for microbial genera. Furthermore, we measured microbial diversity using Shannon indices, which varied between gut compartments and dietary treatments. In the midgut, significant differences were noted between the acclimation diet and the other diets (gent/guar, guar, and soya/feather), with these treatments showing higher median diversity than the acclimation condition (Fig. 3b). Overall, the stomach and midgut exhibited higher diversity across all diets compared to the hindgut. In the soya/feather diet, the midgut displayed the highest diversity, while the hindgut consistently had the lowest diversity.

Consistent with these alpha diversity patterns, beta diversity analysis revealed distinct clustering of microbial communities. As shown in Fig. 3c, clustering patterns indicate a separation between the hindgut and the other two compartments (stomach and midgut), highlighting the distinct microbial composition of the hindgut samples.

**Fig. 3.**
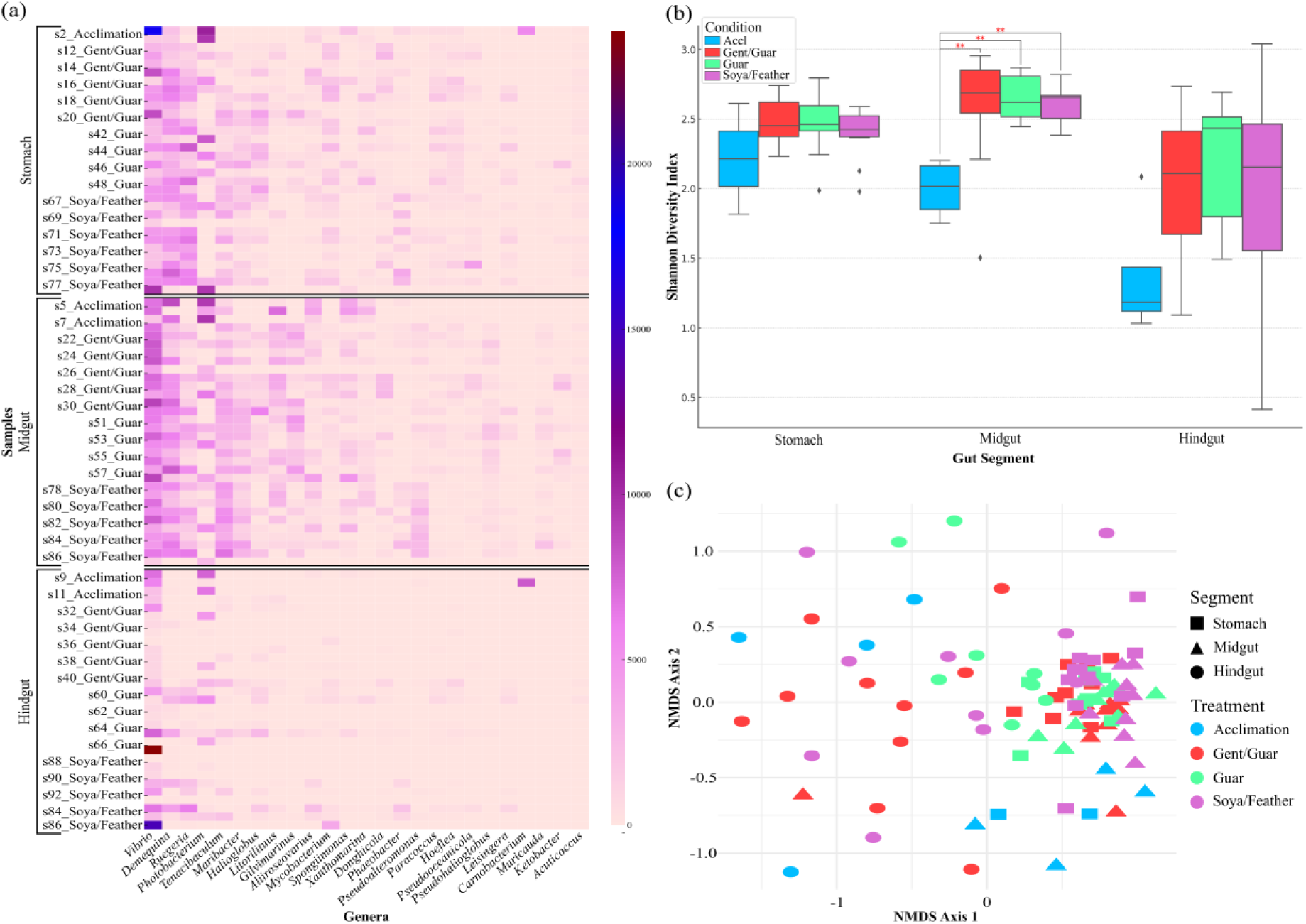
Diet and Compartment-Specific Microbial Diversity and Composition. (a) Heatmap showing the relative abundance of the 25 most abundant genera across samples from stomach, midgut, and hindgut compartments under different diets: acclimation, gent/guar, guar, and soya/feather. Full genus-level abundance data are provided in Supplementary table 1. Genera are displayed on the x-axis, and sample IDs are shown on the y-axis, color-coded by compartment. (b) Shannon diversity indices for stomach, midgut, and hindgut samples reveal variations in microbial diversity across compartments and diets. Boxes represent interquartile ranges, and whiskers indicate data ranges, with significant differences (indicated by asterisks) observed between some diets. (c) Non-metric multidimensional scaling plot of microbial community composition, showing treatment-specific clustering by compartment and diet. Symbols represent compartments (stomach, midgut, hindgut), and colors indicate dietary treatments (acclimation, gent/guar, guar, and soya/feather).

For the gent/guar, guar, and soya/feather diets, samples from the stomach and midgut clustered more closely together, reflecting greater similarity in their microbial communities. In contrast, acclimation samples did not cluster tightly and were more dispersed, suggesting greater variability in microbial communities under this treatment. Building on the observed clustering patterns in the NMDS plot, we performed a PERMANOVA to quantify the variation in microbial composition explained by dietary treatments and gut compartments. The results revealed significant differences in microbial composition across compartments, with the highest variation explained in the midgut, followed by the stomach and hindgut (Fig. 4a). Additionally, a pairwise PERMANOVA (Fig. 4b) provided further insights into the effects of dietary treatments within each compartment. Notably, significant pairwise differences were observed in the midgut and hindgut between specific treatments, such as guar versus soya/feather and acclimation versus gent/guar diet.

**Fig. 4.**
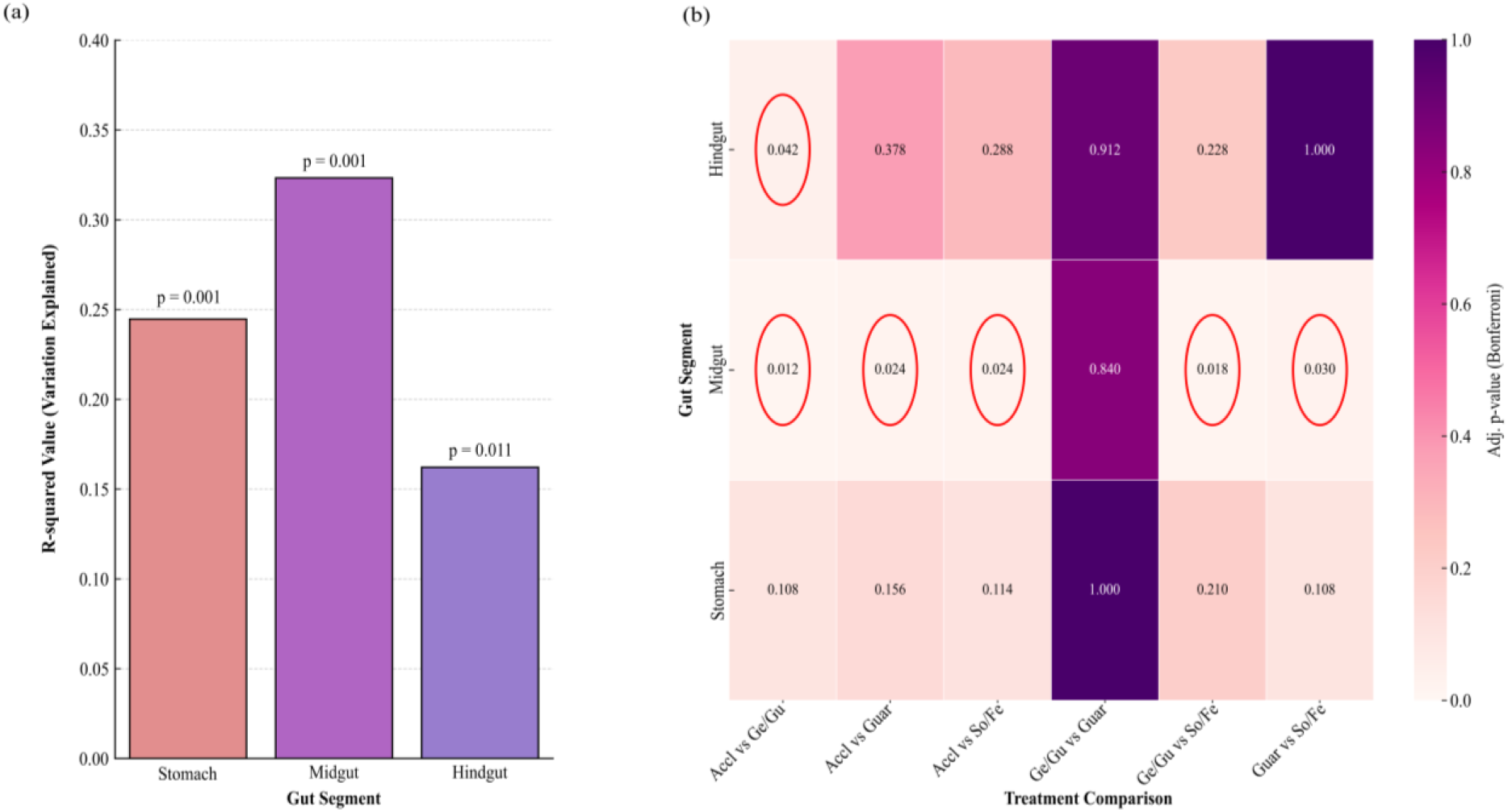
(a) The bar plot shows the variation in microbial composition (R-squared values) explained by gut compartments (stomach, midgut, hindgut) based on PERMANOVA analysis, with associated p-values. The midgut exhibits the highest variation explained (p = 0.001). (b) The heatmap illustrates pairwise adjusted p-values (Bonferroni) from PERMANOVA for different treatments within each compartment. Significant comparisons (adjusted p < 0.05) are highlighted in red circles, indicating specific treatment effects on microbial composition in the midgut and hindgut.

Following the PERMANOVA analysis, we investigated specific bacterial genera contributing to the observed differences in microbial composition. Across all compartments, the acclimation group exhibited the most significant differences in bacterial genera abundance when compared to other treatments (Fig. 5). Most genera that differed significantly due to the transition from acclimation diet to experimental diets showed decreased abundance, as indicated by the predominance of positive log₂ fold changes in these comparisons. Following the transition from acclimation diet to treatment diets (Fig. 5), an exception was observed for *Vagococcus* populations, which were significantly lower due to the experimental diets when compared with the acclimation phase. In the midgut, diet-specific shifts were noted in certain genera; *Amylibacter* abundance increased significantly in the gent/guar compared to the acclimation and guar diets. Conversely, *Marivita* and *Polaribacter* decreased significantly under the guar diet compared to the gent/guar diet.

**Fig. 5.**
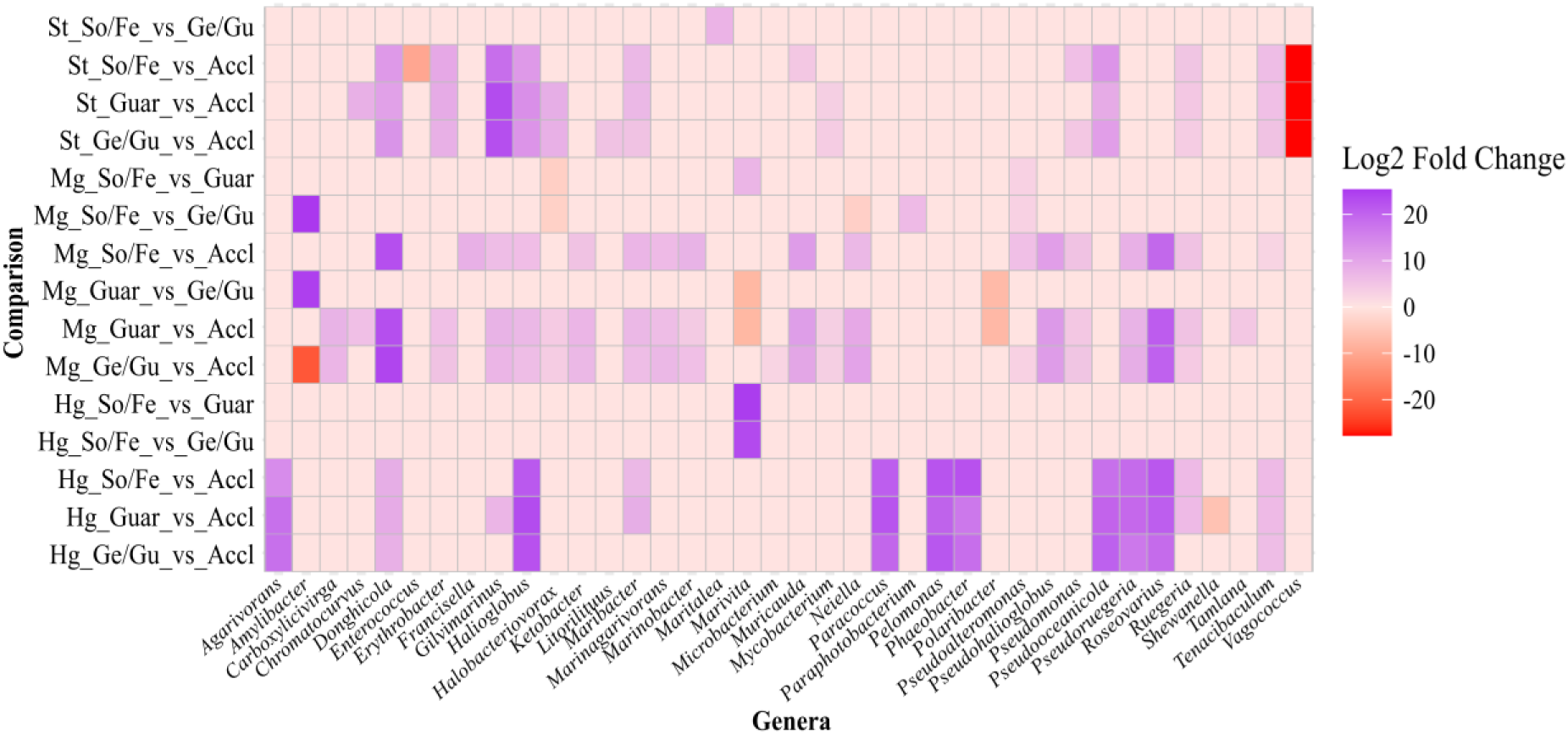
Microbial Genera Shifts Across Diets and Gut Compartments. Heatmap of significant log2 fold changes in genus abundance across treatments within gut compartments. Colors represent the direction and magnitude of fold changes: red indicates negative log2 fold changes (higher abundance in the second condition of each comparison), while purple indicates positive log2 fold changes (higher abundance in the first condition). Cells in misty rose represent either non-significant changes or missing data. Comparisons are arranged by compartment (stomach, midgut, hindgut) along the y-axis, with genera along the x-axis.

Additionally, *Halobacteriovorax* abundance in the midgut increased significantly in the guar and gent/guar diets compared to the acclimation and soya/feather diets; it also increased significantly in the stomach compared to the acclimation group.

To further investigate the microbial taxa driving these differences, we analyzed the abundance of specific bacterial genera across diets and gut compartments (Fig. 6). Remarkably, the acclimation diet did not exhibit any significant microbial changes between compartments, in contrast to the experimental diets, indicating a stable microbial community under control conditions. Conversely, the hindgut displayed the most significant changes in microbial diversity and composition in response to dietary treatments. Under the soya/feather diet, numerous significant shifts were observed between the hindgut and midgut. For instance, the genus *Vibrio* was more abundant in the hindgut compared to the midgut and stomach, reflecting its preferential enrichment in this compartment under the soya/feather diet. Genera such as *Enterococcus* increased significantly in the hindgut of shrimp fed the guar-based and gent/guar diets.

**Fig. 6.**
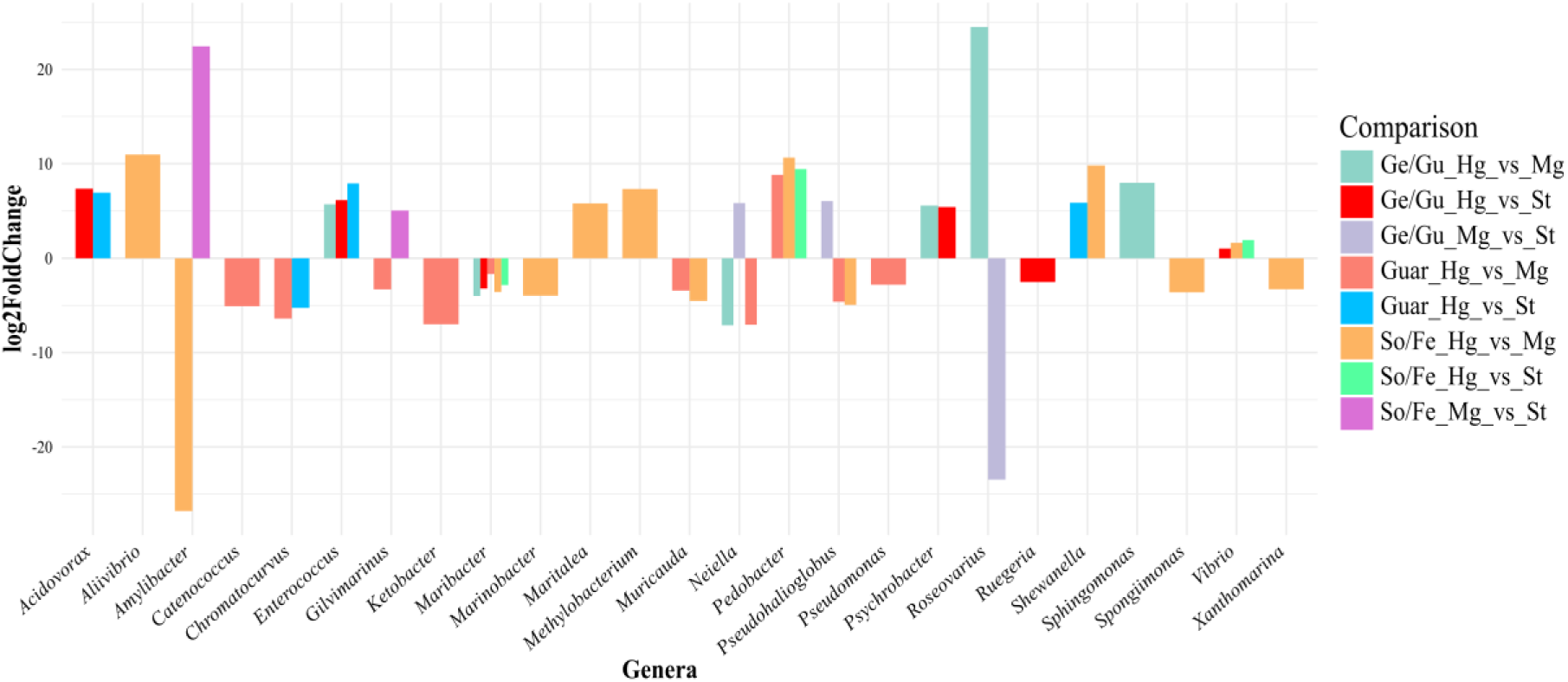
shows significant differential abundance of bacterial genera across shrimp digestive compartments by diet. Positive values indicate higher abundance in the first factor, negative in the second.

Diet- and compartment-specific preferences were evident for other genera as well. *Ruegeria* was less abundant in the hindgut compared to the stomach under the gent/guar diet but showed greater abundance in the hindgut compared to the midgut under the soya/feather diet, suggesting that its distribution is influenced by both diet and gut compartment. Similarly, *Catenococcus* was less abundant in the hindgut compared to the midgut under the guar diet, indicating it thrives more in the midgut under this treatment. *Roseovarius* exhibited notable variations across compartments under the gent/guar diet, being less abundant in the midgut compared to the stomach and more abundant in the hindgut compared to the midgut, highlighting its compartment-specific adaptation. *Pseudomonas* was less abundant in the hindgut compared to the midgut under the guar diet, emphasizing the diet- and compartment-driven microbial dynamics.

## 4. Discussions

As global demand for sustainable aquaculture intensifies, optimizing feed formulations and maintaining efficient digestive function and gut microbiota balance in farmed shrimp such as *Litopenaeus vannamei* is essential for ensuring economic viability and environmental resilience.(61). This study investigates the effects of different dietary formulations, including fiber-rich guar meal, soybean meal, and antibiotic supplementation on digestive enzyme activity, nutrient digestibility, and gut microbiome composition in *L. vannamei*. Our findings reveal that shrimp exhibit specific enzymatic adaptations to varying pH and temperature conditions, with optimal digestive enzyme activities at pH 8 and temperatures ranging between 40°C and 50°C. Additionally, we observed significant diet-induced shifts in nutrient absorption efficiency, particularly higher protein digestibility with guar-based diets compared to soya/feather-based diets. Dynamic microbial responses were noted across gut compartments, with diet-specific alterations in microbial diversity and composition.

### 4.1. Influence of diet on shrimp digestive enzyme activity

The acclimation diet, closely matching a typical aquaculture diet for shrimp, consistently supported higher enzyme activity levels, particularly for leucine aminopeptidase (Fig. 1). The significant reduction in leucine aminopeptidase activity observed in shrimp fed the experimental diets (guar-based, gent/guar, and soya/feather-based) suggests that this enzyme is particularly susceptible to dietary composition (62), and possibly the antinutritional factors present in soy meal and guar meal. In contrast, amylase activity remained stable across different diets, reinforcing its essential role in carbohydrate metabolism (63, 64) regardless of dietary changes. Furthermore, the composition of the experimental diets provides critical insights into these observations. The guar-based diet contained 20% guar meal, which is rich in soluble fiber that increases intestinal viscosity (65, 66). Increased viscosity can reduce the efficiency of enzyme-substrate interactions (67, 68), potentially leading to decreased leucine aminopeptidase activity compared to the acclimation diet. Despite this reduction in enzyme activity, shrimp fed the guar-based diet exhibited high ADCs for protein, exceeding 90% (Fig. 2). One possible explanation is that soluble fiber in the guar meal influenced digestive dynamics such as gut motility or nutrient transit time (69). However, because our enzymatic assays measured specific activity in tissue extracts rather than total enzymatic capacity relative to organ size or body weight, further studies would be needed to confirm this mechanism. On the other hand, the soya/feather-based diet, containing 21.53% soybean meal, likely interfered with digestion due to the presence of antinutritional factors (ANFs) such as trypsin inhibitors and lectins (69–71). These compounds are known to inhibit protease activities and disrupt digestive processes (72). The reduced total protease activity and lower ADCs (∼75%) observed in shrimp fed the soya/feather-based diet cannot solely be attributed to the ANFs in soybean meal. The inclusion of 20% feather meal, a keratin-rich protein source, likely contributed to this outcome. Feather meal is known for its low digestibility in aquatic species due to its high keratin content, a structural protein resistant to enzymatic degradation (28, 73). Additionally, shrimp lack keratinase, an enzyme essential for breaking down keratin (29), which may have exacerbated the observed reductions in nutrient absorption. Consequently, the presence of feather meal in the diet may have compounded the inhibitory effects of soybean meal ANFs on protease activity, resulting in the overall lower ADCs. The lower inclusion of soy protein concentrate (SPC) in the Soy-based diet (8.55%) compared to the Guar-based diet (16.47%) may have further contributed to reduced protein digestibility, as SPC undergoes additional processing to remove ANFs, enhancing its nutritional quality.

Leucine aminopeptidase activity was significantly higher in the gent/guar diet group compared to the guar-based diet (p = 0.0004). Moreover, the soya/feather-based diet showed significantly higher leucine aminopeptidase activity than the guar-based diet (p = 0.003, Fig. 1). However, no significant difference was found between the gent/guar diet and the soya/feather-based diet. This pattern suggests that the combined effects of dietary fiber in the guar-based diet and ANFs in the soya/feather-based diet differentially influence enzyme function. Gentamicin significantly enhanced leucine aminopeptidase activity compared to the guar diet, potentially by modulating the gut microbiota in a way that reduced microbial competition for host nutrients or lowered inflammation, thereby improving epithelial enzyme expression. This effect is consistent with previous studies in poultry, where antibiotics have been shown to enhance digestive enzyme activity and nutrient utilization by suppressing subclinical gut disturbances (74). While leucine aminopeptidase activity was significantly affected by diet, amylase, protease and trypsin activities showed no significant differences across diets (Fig. 1). This stability may simply reflect that the dietary treatments used in this study did not substantially affect hepatopancreatic enzyme secretion.

Differences in diet composition, particularly in protein sources, further explain the observed effects. The higher SPC content in the guar-based diet may have provided a more digestible protein source, mitigating the negative effects of soluble fiber. Both diets included lysine and methionine supplementation to balance essential amino acids. However, imbalances may still occur, particularly since the formulations were originally designed for rainbow trout, whose EAA requirements differ from those of *L. vannamei*. For instance, shrimp have relatively higher requirements for methionine and arginine, while trout require more lysine and leucine for optimal growth and protein deposition (75–77). Such differences may influence protein utilization efficiency and contribute to the variation observed in ADCs. The significantly lower digestibility of the soya/feather-based diet compared to the guar-based and gent/guar diets may be influenced by residual antinutritional factors (ANFs) in soybean meal. However, as protease activity was not significantly affected (Fig. 1), and the diet underwent extrusion, a process known to denature protein-based ANFs (78). Other factors such as the inclusion of poorly digestible feather meal likely played a more dominant role.

The lower SPC content in the soya/feather-based diet exacerbates this issue, as SPC provides a more digestible protein source with reduced ANFs. Additionally, soybean meal has been documented to negatively affect gut health and microbial balance, potentially promoting dysbiosis (79–81), as indicated by the altered microbial diversity observed in our study.

What’s more, the addition of gentamicin did not significantly affect apparent digestibility between the guar-based and gent/guar diets (Fig. 2). While antibiotics are sometimes used as growth promoters to improve nutrient digestibility in livestock by modifying gut microflora (82), their impact can vary across species and contexts. In aquaculture, antibiotic use can disrupt the gut microbiota, potentially impairing digestive processes (83).

### 4.2. Dietary influence on gut microbial composition and diversity

The gut microbiota of *Litopenaeus vannamei* plays a pivotal role in maintaining host physiological homeostasis, contributing to nutrient digestion, immune function, and overall health (84). Our research builds on previous studies by examining microbial communities across all major sections of the digestive tract in crustaceans (37, 85), including the foregut (stomach), midgut (hepatopancreas) and hindgut (intestine), providing further insight into their microbiota. The findings indicate that dietary composition significantly influences the gut microbiota’s diversity and composition, with pronounced effects observed in shrimp fed experimental diets compared to those on the acclimation diet. The experimental diets, guar-based, gent/guar, and soya/feather-based, introduced dietary components such as soluble fiber from guar meal and proteins from soybean meal. These components likely served as substrates for specific microbial populations, altering the gut environment and promoting the growth of polysaccharide and protein-degrading microbes, thereby increasing microbial diversity.

In the midgut, no significant differences in alpha and beta diversity were observed between shrimp fed the guar-based diet and those receiving the same diet supplemented with gentamicin (Fig. 3 b,c). This suggests that the addition of gentamicin did not substantially alter the microbial community established by the fiber-rich guar diet. Dietary fiber is known to modulate gut microbiota composition by promoting beneficial microbial populations (86), which may enhance resilience against antibiotics. Specific microbial taxa responded differently to dietary treatments. For instance, *Amylibacter* decreased significantly in the midgut of shrimp fed the gent/guar diet compared to those on the guar-based diet and acclimation (Fig. 5). The experimental diets without gentamicin, such as guar and soya/feather, may have provided substrates that supported its persistence, whereas gentamicin disrupted its niche by altering microbial competition or directly inhibiting its growth (87, 88). In the midgut, *Marivita* and *Polaribacter* decreased significantly under the guar diet compared to the gent/guar diet. This reduction may reflect competition from fiber-degrading microbes enriched by the guar diet, which could have outcompeted *Marivita* and *Polaribacter* for resources. In addition, gentamicin in the gent/guar diet may have reduced competing microbial populations, indirectly supporting the persistence of *Marivita* and *Polaribacter* in this treatment. Overall, these results emphasize the profound impact of experimental diets on microbial dynamics, with specific genera responding to the introduction of fiber, protein, and antibiotics in distinct ways. The acclimation diet’s stability contrasts with the experimental diets’ capacity to alter microbial niches, driving significant shifts in genera across compartments. The fiber-rich guar diet significantly decreased *Shewanella* in the hindgut compared to acclimation, possibly due to competition from fiber-adapted microbes, altering fermentation dynamics and microbial community structure. Additionally, the significant increase in *Vagococcus* under the acclimation diet compared to the experimental diets (Fig. 5) further indicates that certain bacteria thrive in less perturbed gut conditions.

Our results also highlight the compartmental specialization within the shrimp gut. What stands out first is that the acclimation diet did not exhibit any significant microbial changes between compartments, in contrast to the experimental diets. This stability likely reflects the acclimation diet’s alignment with the shrimp’s natural nutritional requirements, maintaining a balanced and homeostatic microbial ecosystem across the digestive tract. The absence of modulating factors, such as fiber from guar, protein variability from soya/feather, or antibiotic intervention from gentamicin, suggests that the acclimation diet supports uniform microbial populations across compartments, minimizing compartment-specific microbial shifts. The hindgut exhibited the most significant changes in microbial diversity and composition in response to dietary treatments, reflecting its function as the primary site of microbial fermentation (37, 89). The midgut, involved in nutrient absorption (90, 91), showed diet-induced shifts but to a lesser extent. In contrast, the stomach, characterized by a transient and acidic environment (92, 93), displayed reductions in bacterial genera across treatments, suggesting reduce microbial colonization in this compartment.

Genera such as *Enterococcus* increased significantly in the hindgut of shrimp fed the guar-based and gent/guar diets. The hindgut’s role as a fermentation chamber (37), coupled with the availability of soluble fiber, likely created favorable conditions for lactic acid bacteria like *Enterococcus*, which contribute to gut health and nutrient metabolism (94, 95). The resilience of *Enterococcus* to gentamicin suggests its adaptability and potential probiotic role under fiber-enriched conditions (96).

The soya/feather-treated group exhibited numerous significant shifts between the hindgut and midgut, reflecting the pronounced impact of soybean meal on compartment-specific microbial dynamics. The soya/feather diet, rich in protein and might contained antinutritional factors such as trypsin inhibitors and lectins (18, 33), likely disrupted microbial stability, driving divergence between the midgut and hindgut communities. The interplay between diet, gut microbiota, and digestive enzyme activity is evident when considering the impact of dietary components on both microbial communities and enzyme function. The soluble fiber in the guar-based diet not only influenced microbial diversity but also affected digestive enzyme activity, as discussed earlier. The enrichment of fiber-degrading microbes in the hindgut may enhance the breakdown of complex carbohydrates, potentially compensating for reduced enzymatic activity in the shrimp’s digestive tract. Lastly, the lack of significant differences in microbial diversity between the guar-based and gent/guar diets suggests that dietary fiber may confer resilience against antibiotic-induced disruptions. This finding underscores the potential of dietary interventions to modulate the gut microbiota and mitigate the negative effects of antibiotics, which is particularly relevant given the concerns over antibiotic use in aquaculture (97).

Overall, our study demonstrates that dietary composition profoundly influences the gut microbiota of *L. vannamei*, with potential implications for shrimp nutrition and digestive function. The introduction of soluble fiber and ANFs through experimental diets altered microbial diversity and composition, particularly in gut compartments involved in digestion and fermentation. Understanding these interactions is an important step toward developing dietary strategies that support efficient nutrient utilization and may help maintain a balanced gut microbiota conducive to overall shrimp performance.

## 5. Conclusions

The results of this study highlight the complex interactions between dietary composition, digestive enzyme activity, and the gut microbiota in *L. vannamei*. Our findings suggest that guar-based diets, despite containing soluble fiber, supported high protein digestibility and relatively stable microbial communities, making them promising candidates for future feed formulations. In contrast, the soya/feather-based diet resulted in lower protein digestibility and more pronounced microbial shifts, likely due to a combination of less digestible protein sources and residual antinutritional factors. While the addition of gentamicin slightly altered microbial composition and increased leucine aminopeptidase activity, it did not improve digestibility beyond the guar-based diet alone, suggesting limited added value under these conditions. Moving forward, shrimp nutrition research should further evaluate fiber-rich ingredients such as guar meal, in combination with highly digestible protein sources like soy protein concentrate and focus on non-antibiotic strategies to modulate gut microbiota. A more integrative approach, considering enzyme function, microbiota composition, and ingredient digestibility, can help develop cost-effective, high-performing diets that reduce reliance on fishmeal and antibiotics, advancing a more sustainable model of shrimp aquaculture.

## 7. Ethics declarations

This study was approved by the School of Veterinary Medicine Research Ethics Committee, University of Glasow.

## 9. Data availability statement

Raw sequencing data (16S rRNA gene amplicons) have been deposited in the NCBI Sequence Read Archive (SRA) under BioProject accession number PRJNA1286080.

## References

1. Strebel LM, Nguyen K, Araujo A, Corby T, Rhodes M, Beck BH, et al. On demand feeding and the response of Pacific white shrimp (Litopenaeus vannamei) to varying dietary protein levels in semi-intensive pond production. Aquaculture. 2023;574:739698.

2. Liu A, Dumas A, Hernandez JM, Santigosa E. Vitamin nutrition in shrimp aquaculture: A review focusing on the last decade. Aquaculture. 2024;578:740004.

3. Eggink KM, Gonçalves R, Skov PV. Shrimp processing waste in aquaculture feed: nutritional value, applications, challenges, and prospects. Reviews in Aquaculture. 2024.

4. Jastaniah SD, Alaidaroos BA, Shafi ME, Aljarari RM, Abd El-Aziz YM, Munir MB, et al. Dietary Pediococcus acidilactici improved the growth performance, feed utilization, gut microbiota, and disease resistance against Fusarium solani in Pacific white shrimp, Litopenaeus vannamei. Aquaculture International. 2024;32(3):3195–215.

5. Lu M, Liu R, Chen Z, Su C, Pan L. Effects of dietary dihydromyricetin on growth performance, antioxidant capacity, immune response and intestinal microbiota of shrimp (Litopenaeus vannamei). Fish & Shellfish Immunology. 2023;142:109086.

6. Liu Q, Gao Y, Wang C, Zeng Y, Lai Z. Proportions of Pacific white shrimp, Litopenaeus vannamei, gut microbiota from ambient microbiota increased with aquaculture process. Journal of the World Aquaculture Society. 2023;54(4):982–93.

7. Ma Q, Zhao G, Liu J, Chen I-T, Wei Y, Liang M, et al. Effects of a phytobiotic-based additive on the growth, hepatopancreas health, intestinal microbiota, and Vibrio parahaemolyticus resistance of Pacific white shrimp, Litopenaeus vannamei. Frontiers in Immunology. 2024;15:1368444.

8. Mendoza-Porras O, Broadbent JA, Beale DJ, Escobar-Correas SM, Osborne SA, Simon CJ, Wade NM. Post-prandial response in hepatopancreas and haemolymph of Penaeus monodon fed different diets. Omics insights into glycoconjugate metabolism, energy utilisation, chitin biosynthesis, immune function, and autophagy. Comparative Biochemistry and Physiology Part D: Genomics and Proteomics. 2023;46:101073.

9. Fu Q, Sun K, Sui J, Li X, Cao J, Tan J, et al. Comparisons and genetic assessments of WSSV resistance and growth in strain cross of Litopenaeus vannamei. Aquaculture Reports. 2023;30:101572.

10. Li Z, Ju C, Jiao T, Liu H, Li Q. Effects of biofloc on growth performance and survival of Pacific white shrimp (Litopenaeus vannamei) through C/N ratio manipulation, probiotic supplementation, and cocultivation time: A meta-analysis. Aquaculture. 2024:740837.

11. Raza B, Zheng Z, Zhu J, Yang W. A Review: Microbes and Their Effect on Growth Performance of Litopenaeus vannamei (White Leg Shrimps) during Culture in Biofloc Technology System. Microorganisms. 2024;12(5):1013.

12. Emerenciano MG, Rombenso AN, Vieira FdN, Martins MA, Coman GJ, Truong HH, et al. Intensification of penaeid shrimp culture: an applied review of advances in production systems, nutrition and breeding. Animals. 2022;12(3):236.

13. Abualreesh MH. Biodiversity and contribution of natural foods in tiger shrimp (Penaeus monodon) aquaculture pond system: a review. 2021.

14. Nunes AJ, Dalen LL, Leonardi G, Burri L. Developing sustainable, cost-effective and high-performance shrimp feed formulations containing low fish meal levels. Aquaculture reports. 2022;27:101422.

15. Chen Y, Mitra A, Rahimnejad S, Chi S, Kumar V, Tan B, et al. Retrospect of fish meal substitution in Pacific white shrimp (Litopenaeus vannamei) feed: Alternatives, limitations and future prospects. Reviews in Aquaculture. 2024;16(1):382–409.

16. Boyd CE, McNevin AA, Davis RP. The contribution of fisheries and aquaculture to the global protein supply. Food security. 2022;14(3):805–27.

17. Macusi ED, Liguez CGO, Macusi ES, Liguez AKO, Digal LN. Factors that influence small-scale Fishers’ readiness to exit a declining fishery in Davao Gulf, Philippines. Ocean & Coastal Management. 2022;230:106378.

18. Hua K, Cobcroft JM, Cole A, Condon K, Jerry DR, Mangott A, et al. The future of aquatic protein: implications for protein sources in aquaculture diets. One Earth. 2019;1(3):316–29.

19. Kari ZA, Sukri SAM, Rusli ND, Mat K, Mahmud M, Zakaria NNA, et al. Recent advances, challenges, opportunities, product development and sustainability of main agricultural wastes for the aquaculture feed industry–a review. Annals of Animal Science. 2023;23(1):25–38.

20. Amiin MK, Lahay AF, Putriani RB, Reza M, Putri SME, Sumon MAA, et al. The role of probiotics in vannamei shrimp aquaculture performance–A review. Veterinary World. 2023;16(3):638.

21. Zhou L, Han F, Lu K, Qiao Y, Li E. Comparative study on prebiotic effects of different types of prebiotics in an in vitro fermentation by gut microbiota of shrimp (Litopenaeus vannamei). Aquaculture. 2023;574:739687.

22. Ballantyne R, Lee J-W, Wang S-T, Lin J-S, Tseng D-Y, Liao Y-C, et al. Dietary administration of a postbiotic, heat-killed Pediococcus pentosaceus PP4012 enhances growth performance, immune response and modulates intestinal microbiota of white shrimp, Penaeus vannamei. Fish & Shellfish Immunology. 2023;139:108882.

23. Tao L-t, Lu H, Xiong J, Zhang L, Sun W-w, Shan X-f. The application and potential of postbiotics as sustainable feed additives in aquaculture. Aquaculture. 2024:741237.

24. Hussain SM, Bano AA, Ali S, Rizwan M, Adrees M, Zahoor AF, et al. Substitution of fishmeal: Highlights of potential plant protein sources for aquaculture sustainability. Heliyon. 2024.

25. Roncarati A, Galosi L, Di Cerbo A, Quagliardi M, Marchetti F, Fiordelmondo E, et al. Effect of a Guar Meal Protein Concentrate in Replacement of Conventional Feedstuffs on Productive Performances and Gut Health of Rainbow Trout (Oncorhynchus mykiss). Fishes. 2024;9(8):295.

26. Dileep N, Pradhan C, Peter N, Kaippilly D, Sashidharan A, Sankar T. Nutritive value of guar and copra meal after fermentation with yeast Saccharomyces cerevisiae in the diet of Nile tilapia, Oreochromis niloticus. Tropical Animal Health and Production. 2021;53:1–13.

27. Macusi ED, Cayacay MA, Borazon EQ, Sales AC, Habib A, Fadli N, Santos MD. Protein fishmeal replacement in aquaculture: A systematic review and implications on growth and adoption viability. Sustainability. 2023;15(16):12500.

28. Shahabuddin A, Al-Munim S, Hossain MF, Anny SA, Hemal S, Rifat MAR, Sutrodhar S. Feather meal as a sustainable protein source for aquaculture in Bangladesh: Economic implications. Journal of Aquatic Research and Sustainability. 2024;1(01):21–8.

29. Brandelli A, Daroit DJ, Riffel A. Biochemical features of microbial keratinases and their production and applications. Applied microbiology and biotechnology. 2010;85:1735–50.

30. Suloma A, El-Husseiny O, Hassane M, Mabroke R, El-Haroun E. Complementary responses between hydrolyzed feather meal, fish meal and soybean meal without amino acid supplementation in Nile tilapia Oreochromis niloticus diets. Aquaculture international. 2014;22:1377–90.

31. Fornari DC, Nazeer S, Weldon A, Davis DA. The efficacy of hydrolized feathermeal as a protein source in diets for juvenile catfish Ictalurus punctatus. Aquaculture. 2023;576:739823.

32. Dios D. Fishmeal replacement with feather-enzymatic hydrolyzates co-extruded with soya-bean meal in practical diets for the Pacific white shrimp (Litopenaeus vannamei). Aquaculture Nutrition. 2001;7(3).

33. Asghar MU, Sajid QUA, Wilk M, Konkol D, Korczyński M. Influence of various methods of processing soybeans on protein digestibility and reduction of nitrogen deposits in the natural environment–a review. Annals of Animal Science. 2024.

34. Bae J, Hamidoghli A, Djaballah MS, Maamri S, Hamdi A, Souffi I, et al. Effects of three different dietary plant protein sources as fishmeal replacers in juvenile whiteleg shrimp, Litopenaeus vannamei. Fisheries and Aquatic Sciences. 2020;23:1–6.

35. Abd El-Naby AS, Eid A, Gaafar AY, Sharawy Z, Khattaby A, El-sharawy MS, El Asely AM. Overall evaluation of the replacement of fermented soybean to fish meal in juvenile white shrimp, Litopenaeus vannamei diet: growth, health status, and hepatopancreas histomorphology. Aquaculture International. 2024;32(2):1665–83.

36. Yun H, Shahkar E, Hamidoghli A, Lee S, Won S, Bai SC. Evaluation of dietary soybean meal as fish meal replacer for juvenile whiteleg shrimp, Litopenaeus vannamei reared in biofloc system. International Aquatic Research. 2017;9(1):11–24.

37. Garibay-Valdez E, Cicala F, Martinez-Porchas M, Gómez-Reyes R, Vargas-Albores F, Gollas-Galván T, et al. Longitudinal variations in the gastrointestinal microbiome of the white shrimp, Litopenaeus vannamei. PeerJ. 2021;9:e11827.

38. Guéganton M, Rouxel O, Durand L, Cueff-Gauchard V, Gayet N, Pradillon F, Cambon-Bonavita M-A. Anatomy and symbiosis of the digestive system of the vent shrimp rimicaris exoculata and rimicaris chacei revealed through imaging approaches. Frontiers in Marine Science. 2022;9:903748.

39. Chaudhary DK, Kim S-E, Park H-J, Kim K-H. Unveiling the Bacterial Community across the Stomach, Hepatopancreas, Anterior Intestine, and Posterior Intestine of Pacific Whiteleg Shrimp. Journal of Microbiology and Biotechnology. 2024;34(6):1260.

40. Zhang R, Shi X, Liu Z, Sun J, Sun T, Lei M. Histological, physiological and transcriptomic analysis reveal the acute alkalinity stress of the gill and hepatopancreas of Litopenaeus vannamei. Marine biotechnology. 2023;25(4):588–602.

41. Ceccaldi H, editor Anatomy and physiology of digestive tract of Crustaceans Decapods reared in aquaculture. Advances in Tropical Aquaculture, Workshop at Tahiti, French Polynesia, 20 Feb-4 Mar 1989; 1989.

42. Ren X, Wang Q, Shao H, Xu Y, Liu P, Li J. Effects of low temperature on shrimp and crab physiology, behavior, and growth: a review. Frontiers in Marine Science. 2021;8:746177.

43. Seethalakshmi P, Rajeev R, Kiran GS, Selvin J. Shrimp disease management for sustainable aquaculture: innovations from nanotechnology and biotechnology. Aquaculture International. 2021;29:1591–620.

44. Harpeni E, Isnansetyo A, Istiqomah I, Murwantoko. Bacterial biocontrol of vibriosis in shrimp: A review. Aquaculture International. 2024:1–31.

45. Prabina D, Swaminathan TR, Mohandas SP, Anjana J, Manjusha K, Preena P. Investigation of antibiotic-resistant vibrios associated with shrimp (Penaeus vannamei) farms. Archives of Microbiology. 2023;205(1):41.

46. Okocha RC, Olatoye IO, Adedeji OB. Food safety impacts of antimicrobial use and their residues in aquaculture. Public health reviews. 2018;39:1–22.

47. Luu QH, Nguyen TBT, Nguyen TLA, Do TTT, Dao THT, Padungtod P. Antibiotics use in fish and shrimp farms in Vietnam. Aquaculture Reports. 2021;20:100711.

48. Marimuthu V, Shanmugam S, Sarawagi AD, Kumar A, Kim IH, Balasubramanian B. A glimpse on influences of feed additives in aquaculture. EFood. 2022;3(1-2):e6.

49. Baltierra-Trejo E, Márquez-Benavides L, Sánchez-Yáñez JM. Inconsistencies and ambiguities in calculating enzyme activity: The case of laccase. Journal of microbiological methods. 2015;119:126–31.

50. Durai V, Lloyd Chrispin C, Bharathi S, Velselvi R, Karthy A. Factors determining the economic performance of Litopeneaus vannamei (whiteleg shrimp), aquaculture in Tamil Nadu, India. Aquaculture Research. 2022;53(13):4689–96.

51. Erlanger BF, Kokowsky N, Cohen W. The preparation and properties of two new chromogenic substrates of trypsin. Archives of biochemistry and biophysics. 1961;95(2):271–8.

52. Bernfeld P. Methods in enzymology. by SP Colowick and NO Kaplan. Academic Press Inc., New York; 1955.

53. Kazlauskaite R, Cheaib B, Heys C, Ijaz UZ, Connelly S, Sloan W, et al. SalmoSim: the development of a three-compartment in vitro simulator of the Atlantic salmon GI tract and associated microbial communities. Microbiome. 2021;9:1–20.

54. Heys C, Cheaib B, Busetti A, Kazlauskaite R, Maier L, Sloan WT, et al. Neutral processes dominate microbial community assembly in Atlantic salmon, Salmo salar. Applied and Environmental Microbiology. 2020;86(8):e02283–19.

55. Schaal P, Cheaib B, Kaufmann J, Phillips K, Ryder L, McGinnity P, Llewellyn M. Links between host genetics, metabolism, gut microbiome and amoebic gill disease (AGD) in Atlantic salmon. Animal Microbiome. 2022;4(1):53.

56. Kazlauskaite R, Cheaib B, Humble J, Heys C, Ijaz UZ, Connelly S, et al. Deploying an in vitro gut model to assay the impact of the mannan-oligosaccharide prebiotic Bio-Mos on the Atlantic salmon (Salmo salar) gut microbiome. Microbiology Spectrum. 2022;10(3):e01953–21.

57. Hunter JD. Matplotlib: A 2D graphics environment. Computing in science & engineering. 2007;9(03):90–5.

58. Waskom ML. Seaborn: statistical data visualization. Journal of Open Source Software. 2021;6(60):3021.

59. Wickham H. Data Analysis. ggplot2: Elegant Graphics for Data Analysis. Cham: Springer International Publishing; 2016. p. 189–201.

60. Love MI, Huber W, Anders S. Moderated estimation of fold change and dispersion for RNA-seq data with DESeq2. Genome biology. 2014;15:1–21.

61. Muthu CM, Vickram A, Sowndharya BB, Saravanan A, Kamalesh R, Dinakarkumar Y. A comprehensive review on the utilization of probiotics in aquaculture towards sustainable shrimp farming. Fish & Shellfish Immunology. 2024:109459.

62. Li X, Han T, Zheng S, Wu G. Nutrition and functions of amino acids in aquatic crustaceans. amino acids in nutrition and health: Amino acids in the nutrition of companion, zoo and farm animals. 2021:169–98.

63. Jiang Z, Xia S, Zhang D, Liu Q, Xu Y, Wang Y, et al. Effects of Dietary Carbohydrate Sources on Growth Performance, Glucose Metabolism, and Digestive Enzyme Activity of Litopenaeus vannamei. Journal of Shellfish Research. 2022;41(3):399–404.

64. Rodríguez-Viera L, Alpízar D, Mancera J, Perera E. Toward a More Comprehensive View of α-Amylase across Decapods Crustaceans. Biology. 2021;10:947.

65. Gill SK, Rossi M, Bajka B, Whelan K. Dietary fibre in gastrointestinal health and disease. Nature Reviews Gastroenterology & Hepatology. 2021;18(2):101–16.

66. Chen M, Guo L, Nsor-Atindana J, Goff HD, Zhang W, Mao J, Zhong F. The effect of viscous soluble dietary fiber on nutrient digestion and metabolic responses Ⅰ: In vitro digestion process. Food Hydrocolloids. 2020;107:105971.

67. Battilana P, Ornstein K, Minehira K, Schwarz J, Acheson K, Schneiter P, et al. Mechanisms of action of β-glucan in postprandial glucose metabolism in healthy men. European journal of clinical nutrition. 2001;55(5):327–33.

68. Wang M, Xu Z, Sun J, Kim B. Effects of enzyme supplementation on growth, intestinal content viscosity, and digestive enzyme activities in growing pigs fed rough rice-based diet. Asian-Australasian journal of animal sciences. 2008;21(2):270–6.

69. Samtiya M, Aluko RE, Dhewa T. Plant food anti-nutritional factors and their reduction strategies: an overview. Food Production, Processing and Nutrition. 2020;2:1–14.

70. Mukherjee R, Chakraborty R, Dutta A. Role of fermentation in improving nutritional quality of soybean meal—a review. Asian-Australasian journal of animal sciences. 2016;29(11):1523.

71. Gemede HF, Ratta N. Antinutritional factors in plant foods: Potential health benefits and adverse effects. International journal of nutrition and food sciences. 2014;3(4):284–9.

72. Francis G, Makkar HP, Becker K. Antinutritional factors present in plant-derived alternate fish feed ingredients and their effects in fish. Aquaculture. 2001;199(3-4):197–227.

73. Badillo-Zapata D, Vega-Villasante F, Nolasco-Soria H, López-Acuña L, Vargas-Ceballos MA, Barreto-Curiel F. Replacement of fishmeal by hydrolyzed feather meal in diets of juvenile Macrobrachium tenellum (river prawns) and its effect on muscle fatty acids. Latin American Journal of Aquatic Research. 2023;51(5):703–16.

74. Onyeanu CT, Ezenduka EV, Anaga AO. Determination of gentamicin use in poultry farms in Enugu state, Nigeria, and detection of its residue in slaughter commercial broilers. 2020.

75. Wilson R, Cowey C. Amino acid composition of whole body tissue of rainbow trout and Atlantic salmon. Aquaculture. 1985;48(3-4):373–6.

76. Richard L, Blanc P-P, Rigolet V, Kaushik SJ, Geurden I. Maintenance and growth requirements for nitrogen, lysine and methionine and their utilisation efficiencies in juvenile black tiger shrimp, Penaeus monodon, using a factorial approach. British journal of nutrition. 2010;103(7):984–95.

77. Walton M, Cowey C, Coloso RM, Adron J. Dietary requirements of rainbow trout for tryptophan, lysine and arginine determined by growth and biochemical measurements. Fish Physiology and Biochemistry. 1986;2:161–9.

78. Nikmaram N, Leong SY, Koubaa M, Zhu Z, Barba FJ, Greiner R, et al. Effect of extrusion on the anti-nutritional factors of food products: An overview. Food control. 2017;79:62–73.

79. Miao S, Zhao C, Zhu J, Hu J, Dong X, Sun L. Dietary soybean meal affects intestinal homoeostasis by altering the microbiota, morphology and inflammatory cytokine gene expression in northern snakehead. Scientific reports. 2018;8(1):113.

80. Zhang W, Tan B, Deng J, Dong X, Yang Q, Chi S, et al. Mechanisms by which fermented soybean meal and soybean meal induced enteritis in marine fish juvenile pearl gentian grouper. Frontiers in Physiology. 2021;12:646853.

81. Suhr M, Fichtner-Grabowski F-T, Seibel H, Bang C, Franke A, Schulz C, Hornburg SC. Effects of plant-based proteins and handling stress on intestinal mucus microbiota in rainbow trout. Scientific Reports. 2023;13(1):22563.

82. Ronquillo MG, Hernandez JCA. Antibiotic and synthetic growth promoters in animal diets: review of impact and analytical methods. Food control. 2017;72:255–67.

83. Li Z, Lu T, Li M, Mortimer M, Guo L-H. Direct and gut microbiota-mediated toxicities of environmental antibiotics to fish and aquatic invertebrates. Chemosphere. 2023;329:138692.

84. Li E, Xu C, Wang X, Wang S, Zhao Q, Zhang M, et al. Gut microbiota and its modulation for healthy farming of Pacific white shrimp Litopenaeus vannamei. Reviews in Fisheries Science & Aquaculture. 2018;26(3):381–99.

85. Castejón D, Rotllant G, Alba-Tercedor J, Ribes E, Durfort M, Guerao G. Morphological and histological description of the midgut caeca in true crabs (Malacostraca: Decapoda: Brachyura): origin, development and potential role. BMC zoology. 2022;7(1):9.

86. Medawar E, Haange S-B, Rolle-Kampczyk U, Engelmann B, Dietrich A, Thieleking R, et al. Gut microbiota link dietary fiber intake and short-chain fatty acid metabolism with eating behavior. Translational psychiatry. 2021;11(1):500.

87. Hahn FE, Sarre SG. Mechanism of action of gentamicin. The Journal of infectious diseases. 1969;119(4/5):364–9.

88. Karunarathna I, Gunasena P, Gunathilake S, De Alvis K. The clinical use of gentamicin: Indications, mechanism of action, and key considerations. ResearchGate. https://www.researchgate.net/publication/383220508; 2024.

89. Zeng S, He J, Huang Z. The intestine microbiota of shrimp and its impact on cultivation. Applied Microbiology and Biotechnology. 2024;108(1):1–8.

90. Zhou KM, Liu PP, Yao JY, Vasta GR, Wang JX, Wang XW. Shrimp Intestinal Microbiota Homeostasis: Dynamic Interplay Between the Microbiota and Host Immunity. Reviews in Aquaculture. 2024.

91. Sonakowska-Czajka L, Śróbka J, Ostróżka A, Rost-Roszkowska M. Postembryonic development and differentiation of the midgut in the freshwater shrimp Neocaridina davidi (Crustacea, Malacostraca, Decapoda) larvae. Journal of Morphology. 2021;282(1):48–65.

92. Imaizumi K, Tinwongger S, Kondo H, Hirono I. Analysis of microbiota in the stomach and midgut of two penaeid shrimps during probiotic feeding. Scientific Reports. 2021;11(1):9936.

93. Lee Y-K, Lin B-Y, Weng T-H, Huang C-K, Liu C, Liu C-C, et al. Counting and measuring the size and stomach fullness levels for an intelligent shrimp farming system. Connection Science. 2023;35(1):2268878.

94. Sun X, Fang Z, Yu H, Zhao H, Wang Y, Zhou F, et al. Effects of Enterococcus faecium (R8a) on nonspecific immune gene expression, immunity and intestinal flora of giant tiger shrimp (Penaeus monodon). Scientific Reports. 2024;14(1):1823.

95. Chino de la Cruz CM, Cornejo-Granados F, Gallardo-Becerra L, Rodríguez-Alegría ME, Ochoa-Leyva A, López Munguía A. Complete genome sequence and characterization of a novel Enterococcus faecium with probiotic potential isolated from the gut of Litopenaeus vannamei. Microbial Genomics. 2023;9(3):000938.

96. Dias J, Marinho Y, Santos I, Almeida E, Pinheiro S, Silva A, et al. Prospection of Lactobacillus plantarum and Enterococcus faecium with potential species-specific probiotic use in ornamental aquaculture of Betta splendens Regan, 1910. Arquivo Brasileiro de Medicina Veterinária e Zootecnia. 2024;76(03):e13141.

97. Imtiaz N, Anwar Z, Waiho K, Shi C, Mu C, Wang C, Qingyang W. A review on aquaculture adaptation for fish treatment from antibiotic to vaccine prophylaxis. Aquaculture International. 2024;32(3):2643–68.

